# Lessons Learned: Recommendations for Establishing Critical Periodic Scientific Benchmarking

**DOI:** 10.1101/181677

**Authors:** Salvador Capella-Gutierrez, Diana de la Iglesia, Juergen Haas, Analia Lourenco, José María Fernández, Dmitry Repchevsky, Christophe Dessimoz, Torsten Schwede, Cedric Notredame, Josep Ll Gelpi, Alfonso Valencia

## Abstract

The dependence of life scientists on software has steadily grown in recent years. For many tasks, researchers have to decide which of the available bioinformatics software are more suitable for their specific needs. Additionally researchers should be able to objectively select the software that provides the highest accuracy, the best efficiency and the highest level of reproducibility when integrated in their research projects.

Critical benchmarking of bioinformatics methods, tools and web services is therefore an essential community service, as well as a critical component of reproducibility efforts. Unbiased and objective evaluations are challenging to set up and can only be effective when built and implemented around community driven efforts, as demonstrated by the many ongoing community challenges in bioinformatics that followed the success of CASP. Community challenges bring the combined benefits of intense collaboration, transparency and standard harmonization. Only open systems for the continuous evaluation of methods offer a perfect complement to community challenges, offering to larger communities of users that could extend far beyond the community of developers, a window to the developments status that they can use for their specific projects. We understand by continuous evaluation systems as those services which are always available and periodically update their data and/or metrics according to a predefined schedule keeping in mind that the performance has to be always seen in terms of each research domain.

We argue here that technology is now mature to bring community driven benchmarking efforts to a higher level that should allow effective interoperability of benchmarks across related methods. New technological developments allow overcoming the limitations of the first experiences on online benchmarking e.g. EVA. We therefore describe OpenEBench, a novel infra-structure designed to establish a continuous automated benchmarking system for bioinformatics methods, tools and web services.

OpenEBench is being developed so as to cater for the needs of the bioinformatics community, especially software developers who need an objective and quantitative way to inform their decisions as well as the larger community of end-users, in their search for unbiased and up-to-date evaluation of bioinformatics methods. As such OpenEBench should soon become a central place for bioinformatics software developers, community-driven benchmarking initiatives, researchers using bioinformatics methods, and funders interested in the result of methods evaluation.

## Introduction

Scientific research dependence on software, data repositories, and advanced computer science methodologies has dramatically increased over the last years, mostly under the pressure of the increasingly large amounts of data produced by experimental biology. This is especially relevant for many areas of life sciences from omics to live imaging to processing electronic health records. Knowledge can emerge from the analysis of newly generated data, the re-analysis of existing ones, and/or from the combination of both. However, as life sciences data sets become larger, minor differences and inaccuracies in available data and/or used software can have a strong impact on the final results [Marx 2013, Di Tommaso et al 2017]. It is also worth noting that bioinformatics software publications tend to be over-optimistic about the self-assessment of the reported software [Norel et al. 2011, Boulesteix 2015]. Thus, external and independent evaluation of bioinformatics software is needed to overcome such biases. An independent assessment will also assist developers and researchers when selecting the most suitable bioinformatics software for their specific scientific needs.

Critical benchmarking of bioinformatics software adds value to research communities by providing objective metrics in terms of scientific performance, technical reliability, and perceived functionality [Jackson et al. 2011, Friedberg et al. 2015]. At the same time, target criteria agreed within a community are an effective way to stimulate new developments by highlighting challenging areas [Costello and Stolovitzky 2013].

Motivated by the success of CASP (Critical Assessment of Techniques for Protein Structure Prediction) [Moult et al. 1995] and building in the long tradition of benchmarking exercises and competitions in software engineering, a number of similar community-driven benchmark experiments have been organized during the last two decades, for instance: BioCreAtIvE (Critical Assessment of Information Extraction in Biology) [Hirschman et al. 2005], CAFA (Critical Assessment of Functional Annotation) [Radivojac et al. 2013], and QfO (Quest for Orthologs) [Altenhoff et al. 2016]. A more complete list is shown in figure 1 and table 1. Together with their intrinsic scientific impact, these efforts promote the organization of communities around scientific challenges, incentive new methodological developments, and, more importantly, inspire the emergence of new communities in other research areas. However, for the efficient development of bioinformatics methods, tools and web services (referred to as “tools” from here on), continuous automated benchmark services are required to compare the performance of tools on previously agreed data sets and metrics. Continuous benchmarking is even more important for users that tend to have difficulties in choosing the right tool for their research questions, and are not necessarily aware of the latest developments in each of the fields of the bioinformatics methods they need to use. Thus, there are many aspects to consider when organizing a benchmarking effort [Friedberg et al. 2015]. For instance, it is important to define i) roles for participants e.g. whether organizers can actively take part at the challenge as participants; ii) who is running the benchmark infrastructure; iii) how the benchmark is performed: in real-time, online, and/or offline; iv) whether the effort is punctual or continuous in time with periodic updates in data sets and/or evaluated metrics; v) how input data sets are processed. In the case of large data sets, a continuous service will provide more reliable and comprehensive statistics and ranking schemes over time [Eyrich et al. 2003]; vi) the final relevance of the reported indicators and their practical usability by external users, as well as for the design of bioinformatics workflows.

Our focus here is on benchmark efforts which can be automated, and potentially run continuously including the use of new reference data sets as soon as they become available. Such system should be in charge of hosting reference data sets, gathering participants data, measure performance, and produce metrics on-demand. It is important to consider that continuous benchmark services require a stable computational and human infrastructure which may be difficult to fund and maintain over time. CAMEO (Continuous Automated Model EvaluatiOn) [Haas et al. 2013], CompaRNA [Puton et al. 2013], and QfO [Altenhoff et al. 2016] are examples of currently active continuous benchmark initiatives; the first one is focused on the evaluation of protein structure predictions, model quality estimation and contact predictions; the second focuses on the prediction of RNA secondary structure; while the latter one measures orthology predictions from different perspectives, including the use manually curated data as reference. Moreover, several projects have implemented automated and/or continuous benchmark systems in the past, e.g. EVA (EValuation of Automatic protein structure prediction) [Rost and Eyrich 2001, Koh et al. 2003], LiveBench [Bujnicki et al. 2001, Rychlewski and Fischer 2005], and CAFA-SP (Critical Assessment of Fully Automated Structure Prediction) [Fischer et al. 1999], but they have been abandoned in the meantime or integrated into other experiments such as CASP [Moult et al. 1995]. Therefore, we envision that many scientific communities would benefit from a stable, generic and efficient infrastructure devoted to host unattended, periodic and continuous benchmark services.

**Figure 1.**
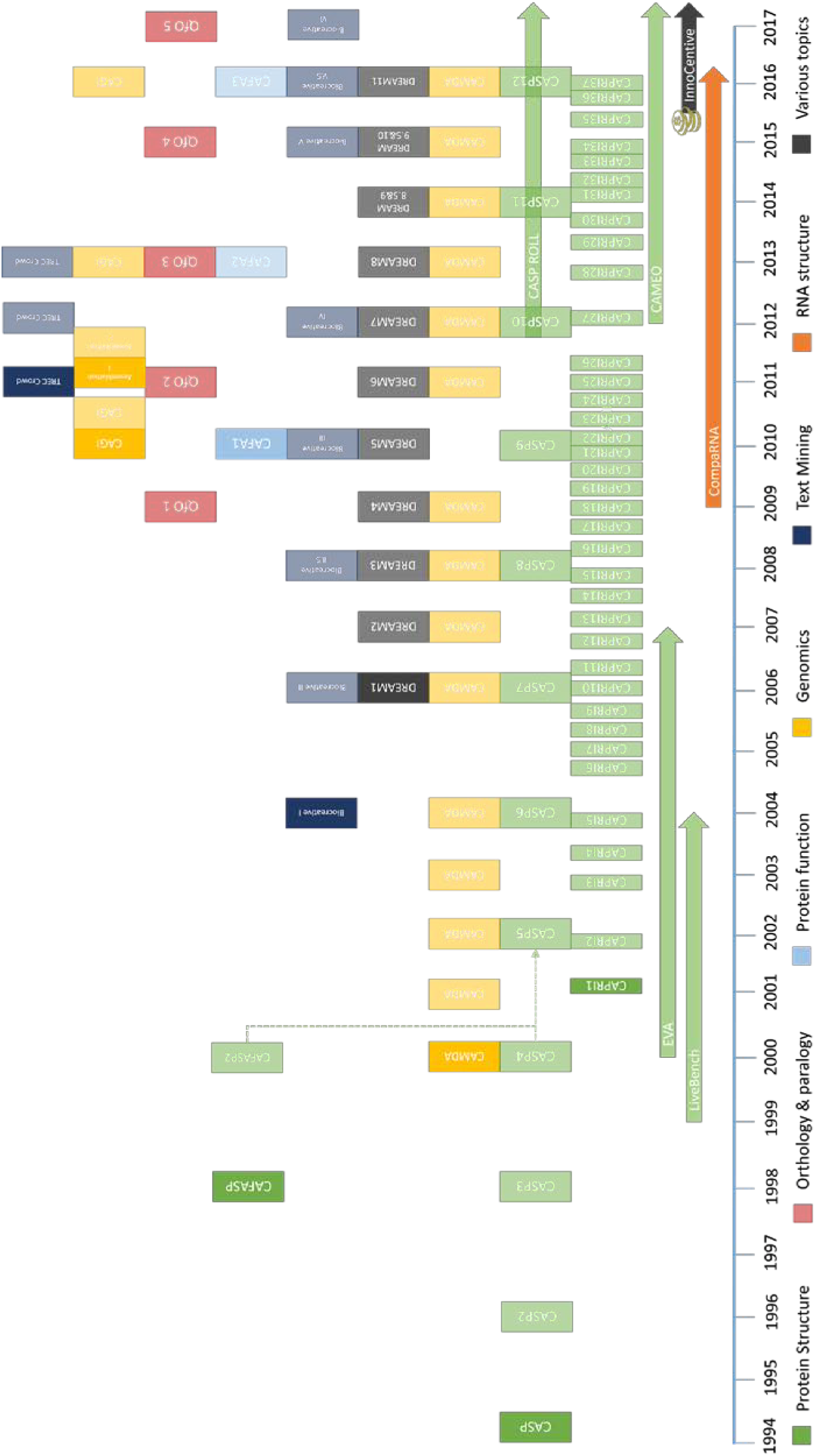
Bioinformatics benchmark initiatives timeline for a broad range of topics. Additional details and other efforts are part of table 1.

A benchmark infrastructure needs a close connection with high-quality repositories for both reference data sets and tools. Despite their primary role in the scientific, technical and/or functional evaluation of tools, such infrastructure should also provide use cases and best practice guidelines about how to establish and maintain communities around benchmark efforts, how to identify scientific relevant questions for each area, and how to make sure data sets and tools implement widely adopted standards for data exchange. The infrastructure should come along with well defined mechanisms allowing its expansion, evolution and long-term sustainability, in tune with the scientific needs of the scientific community.

This work outlines general principles and requirements for designing benchmark initiatives, either continuous or periodic, provides a strengths, weaknesses, opportunities, and threats (SWOT) analysis based on existing communities (Figure 2], and maps active initiatives as example for future efforts. Figure 3 identifies the relevant beneficiaries of community-driven benchmarking initiatives: software developers, communities running benchmark efforts, end-users who benefits of such systems when taking informed decisions about the best choice for their research problem; and funders who can take benefit of an open benchmarking infrastructure to objectively evaluate the contribution of its funding recipients retrospectively and/or prospectively.

**Figure 2.**
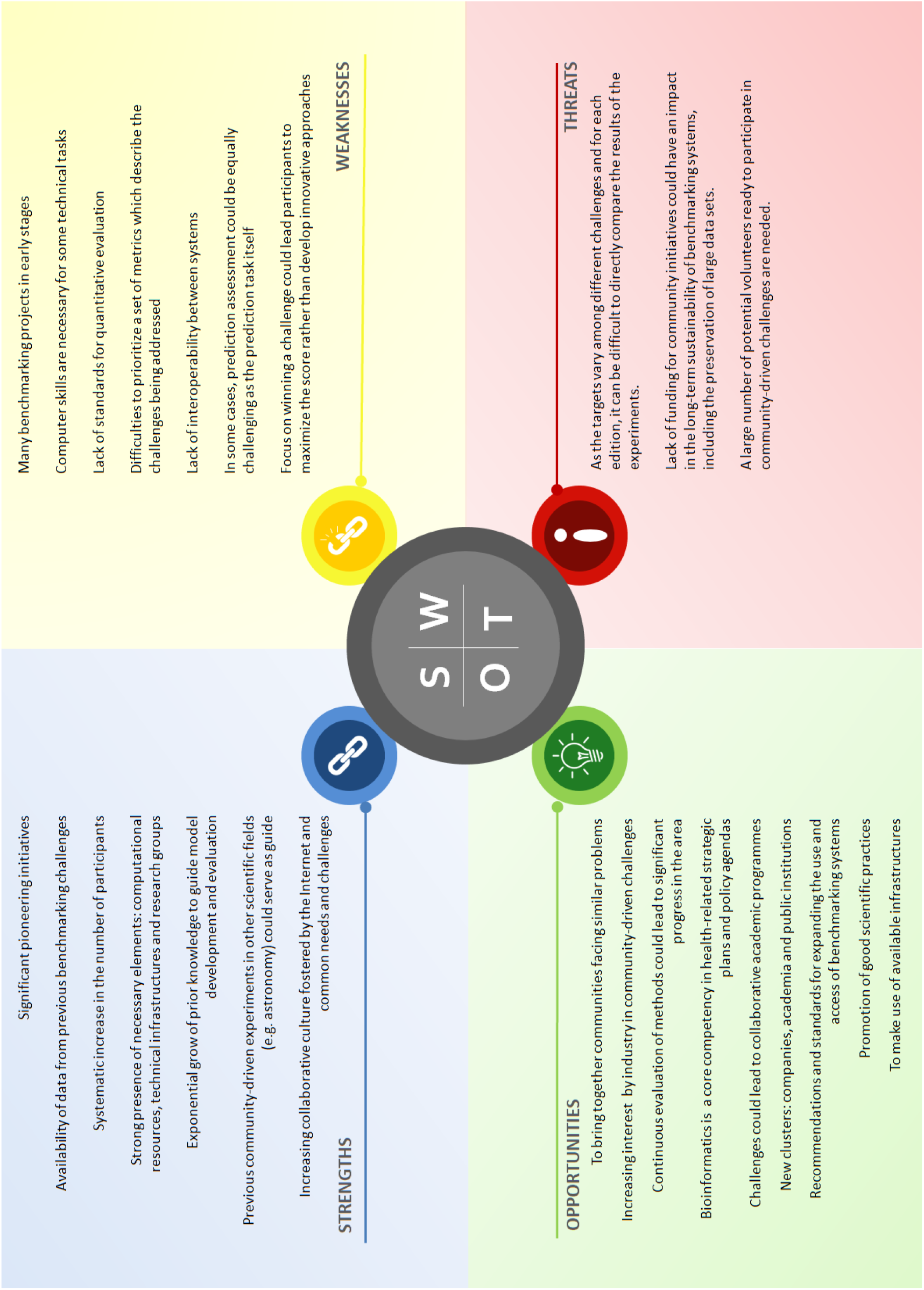
Strengths, weaknesses, opportunities and threats (SWOT) analysis based on existing benchmark initiatives.

**Figure 3.**
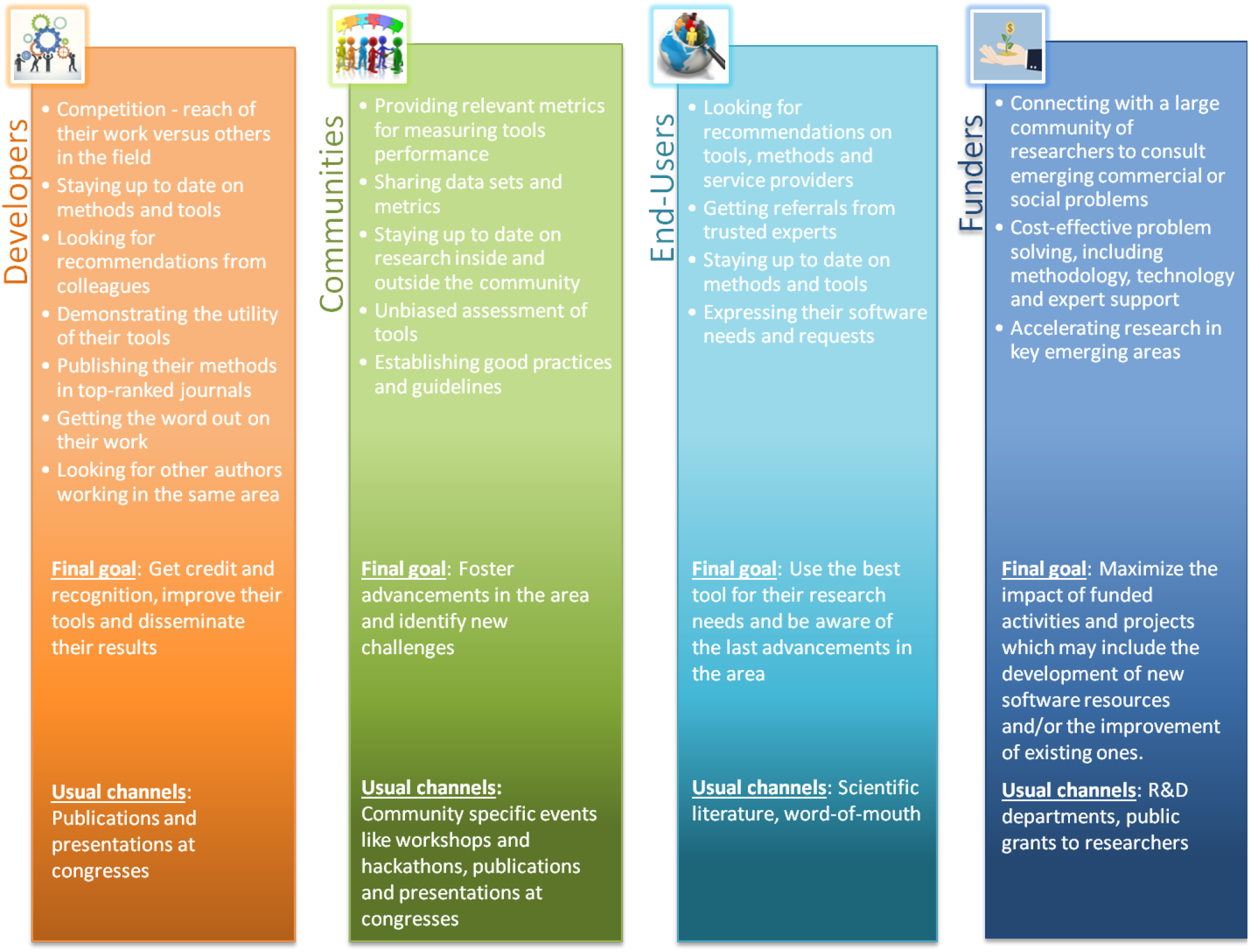
Main actors within a community-driven benchmark challenge: objectives, final goals and previous channels to address their objectives.

## General aspects for establishing a community-driven benchmark effort

Community-driven benchmarking is a complex process entirely relying on intense cooperation among its members. These communities can be effectively strengthened by challenge-based competition with clear participation rules, a scientific sound set of questions, and agreed common data sets (Figure 4]. Provided a critical mass of software developers is reached, competition eventually ends up bringing stimulated rewards and invaluable feedback about potential improvements for individual solutions.

**Figure 4.**
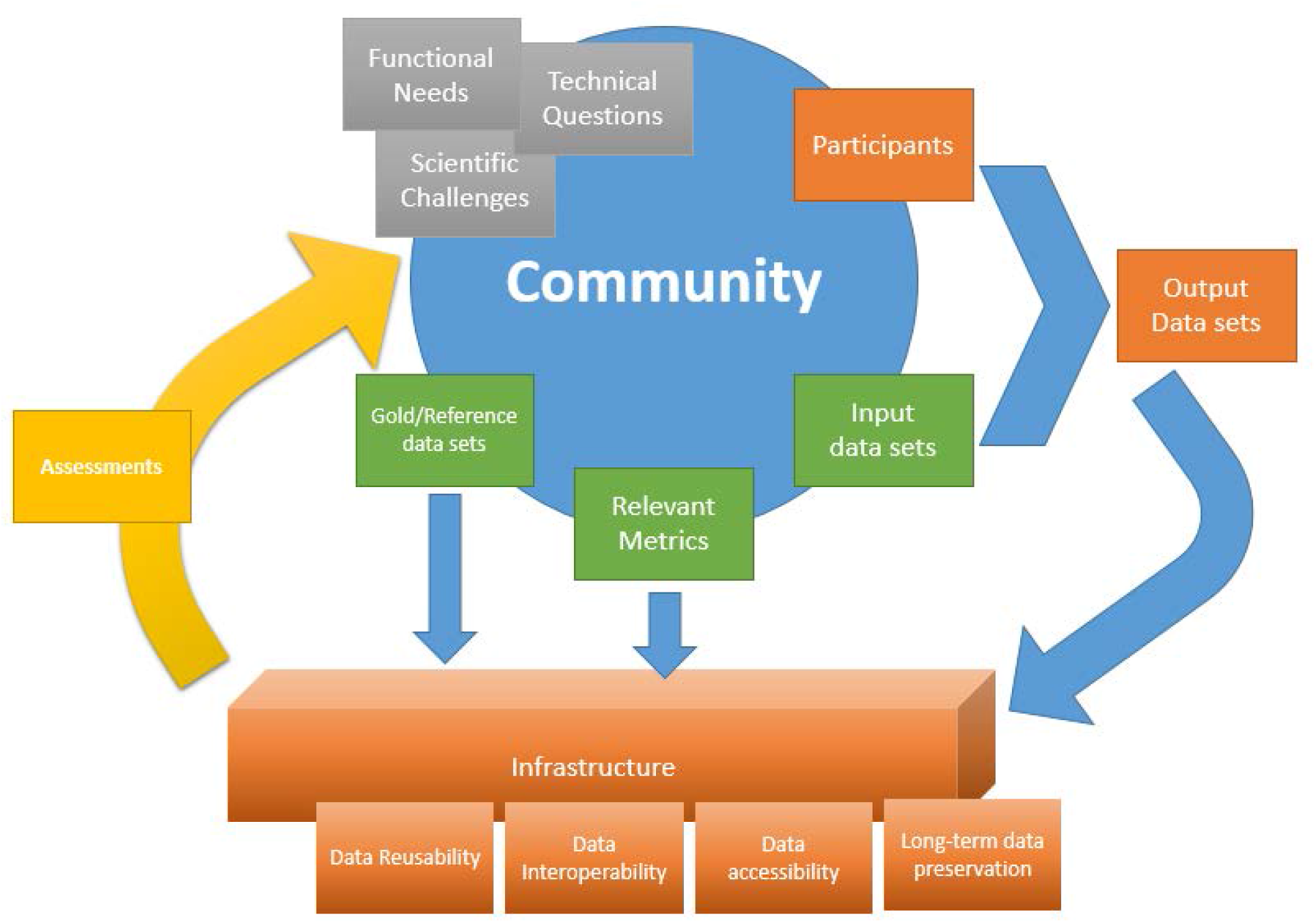
General workflow and crucial input for successful community supported (continuous) evaluations.

The next sections describe several aspects involved in the design of successful challenges which are important and complement previous work in the field [Friedberg et al. 2015].

### 1. Types of benchmarking

General-purpose software is usually evaluated using technical performance metrics, such as average response time and memory access rate. However, in order to add real value for researchers, scientific software cannot be evaluated only from a technical perspective, a more comprehensive evaluation should also measure the scientific performance of software under diverse scenarios of application and data sets.

Evaluation can be carried out by considering three different but yet complementary types of assessment as shown in figure 5.

#### Technical: how does the tool perform under specific technical conditions?

Technical benchmarking usually focuses on elements of technical quality. Relevant factors that determine whether scientific software is technically successful are, for instance, whether it can be compiled with no errors, resources needed along the execution (storage, memory), the reproducibility of the results, and portability, among others. In the case of services, relevant features are accessibility, up-time, communication protocols, response time, processing speed, and interoperability. Importantly, reverse engineering, comparison of technical specifications and analysis of operating statistics are the primary techniques in the context of technical benchmarking.

#### Functional: how usable is a given tool by end-users?

The functional assessment performs a user-based evaluation of software usability. Some relevant aspects that determine the usability of a given software are: how intuitive and easy-to-use is the Graphic User Interface (GUI); if there exists clear and comprehensive user documentation; whether software customizes the user experience according to predefined roles when more than one profile is available; whether it is linked to data repositories that are updated frequently; if there are communities around the software aiming to support users and/or developers; whether the software is open source and licenses are properly indicated.

#### Scientific: how does the method perform within clearly defined scientific challenges?

The main aspect here is to define within the respective community what is to be measured and how, the so-called metrics. These metrics come in all shapes and sizes. Some relate to experimental readouts used as standards of truth while others merely quantify some level of optimization. The metrics serve two complementary purposes: the first one is to objectively evaluate the relative scientific performance of the different participating tools. The second one is more complex and it is related with the understanding of the theoretical basis of the differences between tools, and the evaluation of the specific details of the tools related with their performances. What are the tools potential biases, strength and weaknesses? Under which conditions do tools tend to underperform? Are key scientific questions that cannot be answered easily by the evaluation systems? This is possible since they might require deep scientific knowledge and substantial information about the corresponding tools. However, automated evaluation systems can assist communities to perform this type of analysis by providing the necessary information about the participating tools.

**Figure 5.**
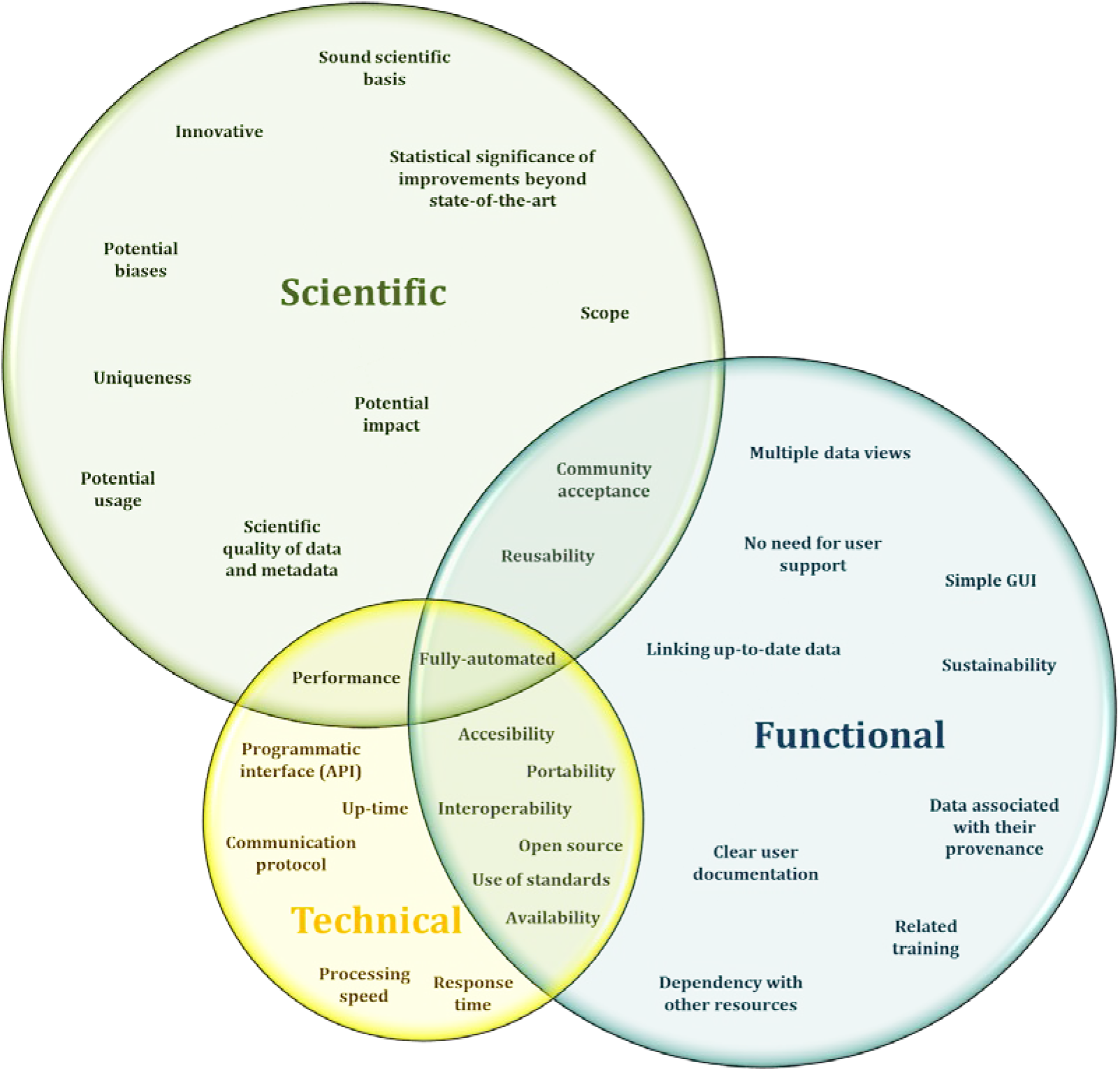
Venn diagram showing relations among the different types of benchmarks: scientific, technical and functional.

### 2. Definition of the benchmark

Understanding why a given software performs as it does on a specific benchmark scenario is often as important as the actual benchmark results [Aniba et al. 2010]. When designing a benchmark, this should be adapted and representative enough of its actual application as well as capable to scaling-up as more participants use it and data volume grows. Moreover, different benchmark editions should move towards more challenging aspects of the core scientific questions under study, so the benchmark design needs to evolve over time to keep its relevance within its scientific domain.

Concise and comprehensive definition of the scientific scope is also key for successfully organizing a benchmarking initiative and attracts participants. The community should define this scope in terms of relevant scientific questions, which can be quantitatively evaluated; and the data needed to answer those questions. Scientific questions should be: achievable, i.e. solvable in a reasonable time, but not trivial; realistic, i.e. representing real problems with real data; and appealing to attract a broad interest. Prediction tasks should be defined in terms of expected results, e.g. range of possible predictions, input and output data formats, and potential impact. Prior to starting the challenge, it is crucial to agree on both the metrics and the evaluation mode, in particular for those cases where a ‘gold standard’ is not available. Importantly, the temporal frame of the challenge should be defined in advance in order to allow the community to learn from it and prepare new editions of the initiative.

Participants and assessors should be involved in the open definition of the scientific challenge and evaluation criteria, and community feedback should be gathered to periodically refine assessment metrics and procedures. Therefore, the evaluation procedure of the benchmark should be transparently accessible to the community at any moment during the challenge implementation. Importantly, benchmark data, including the data sets submitted by participating tools, should be also available after the challenge to facilitate further refinements and the development of new tools using sound and relevant data. Additionally, data considered as reference for the community should be properly identified as such, documented, and maintained at stable repositories.

### 3. Evaluation: reference data sets, metrics and ranking

Continuous benchmarking provides a framework where a large number of tools are being compared and evaluated simultaneously, under the same conditions and over time. As soon as results become available, an assessment on their correctness and/or accuracy has to be carried out. The assessment should allow to effectively compare the performance of participant tools by providing appropriate rankings which are of high interest for developers, the community organized around the benchmark effort, researchers, and funders. The framework must also guarantee the reproducibility of results, and provide ways to verify such reproducibility.

But, which metrics are appropriate to evaluate the accuracy of a result? In scientific benchmarking, a precise definition of what “correct” means is essential. Prediction accuracy can be defined as the degree to which the prediction reflects the real facts of biological systems. However, no single metric can be definitely used to evaluate all the predictions. In almost all cases there are orthogonal aspects necessitating more than one ranking to be made available and, optimally, all scores should be integrated in a way that tool developers and end-users can weight those metrics according to their specific needs. It seems obvious that the benchmark should be method-independent. In the ideal case, a ‘gold standard’, which will be usually kept undisclosed during official challenges, is used to assess the accuracy of predictions applying quantitative metrics, e.g. CAMEO and CASP use new, unpublished, experimental 3D structures that are available on weekly basis. Biocreative uses gold standard manually curated data sets that require laborious and costly production, as well as silver standard data sets which are produced after collecting and combining results from the participating tools. However, this cannot be the case in all communities. Initial efforts around multiple sequence alignments used one of the benchmarked methods output, which was manually refined at a later stage, to build up the reference data sets [Thompson et al. 1999]. The case of multiple sequence alignment (MSA) is especially revealing on the complexity of setting up appropriate gold standards. While MSA modelling is one of the most widely used technique in computational biology [Van Noorden et al., 2014], the agreement across the six main reference datasets has been reported to be as low as 66% [Kemena et al. 2009] with current benchmarks being criticized for deep flaw in their underlying logics. Critics were later raised by other members of the community [Edgar et al. 2010].

Importantly, assessment metrics should be easily interpretable by everyone in the field as this will promote participation. Metrics should also reflect different angles of relevant scientific questions to allow detailed discussions and to avoid convoluted, and, thus, scientifically meaningless scores. Communities, and especially those who volunteered as assessors, should periodically review existing metrics and come up with new metrics reflecting the current developments and challenges in the field.

Communities may use absolute metrics, relative measures and/or statistical significance of differences. Individual metrics are suited to describe different aspects of the evaluation, or the same aspect under different perspectives, and when combined provide a comprehensive view of tools performance. Over time, best methods tend to converge and the relevance of the differences should also be analyzed to determine whether better results are due to the new methods and/or refinement of existing ones, just due to chance, or if they merely reflect an overfitting of the gold standard. Various approaches have been used to evaluate the prediction accuracy of results depending on the availability of such gold standards. For instance, the predictions can be compared directly either with experimental data, with synthetic and/or simulated datasets generated *in silico* following previous experiences [Van den Bulcke et al. 2006, Kim et al. 2010, Hatem et al. 2013] or with data generated using unsupervised learning approaches, based on the consensus among different — i.e. algorithmically independent— methods [Elsik et al. 2007, Chen et al. 2007, Keith et al. 2012]. For the latter, naive methods e.g. Bayesian networks, can provide a baseline allowing assessors to measure relative performance between methods with, on average, moderate to good accuracy. Such consensus data is referred to as ‘silver standard’. However, data from silver standards should be used with caution as it needs to be revised regularly avoiding to poorly evaluate new developments in the field. New developments which implement radical solutions to open scientific questions under scrutiny may produce little to none overlap with existing solutions, and, thus, perform poorly when evaluated with silver standard data sets which are produced after combining the output of existing solutions‥ The use of synthetic data is complicated. On the one hand synthetic data is perfectly suited to any problem dominated by an optimization issue e.g. Maximum Likelihood tree computation; on the other hand synthetic data is poorly suited to be a substitute for experimental data, especially when the biological process is imperfectly modelled.

In order to extract valuable conclusions from the results, sometimes, it is also necessary to use additional information, such as expert analysis and manual curation. Proper ontological description of the references and their associated metrics would therefore be instrumental at ensuring that metrics and datasets are described in the most informative way with respect to the aspect of the modelling they are capturing.

To guarantee a fair comparison, it is essential that participants use only input data agreed by the community. Hosting such data in well defined data repositories will prevent having distorted results because of different IDs for the same input data and/or different data versions. In the case of fully automated approaches, where continuous benchmarking is achieved by interrogating web servers, additional strategies may be necessary. Another important aspect is the amount of available data for conducting benchmarking activities because few cases might lead to similar results among participating tools. In fact, input data sets should contain varied test cases covering a wide range of possible inputs to the prediction problem. This provides means to prove the ability of participants to generalize and conform to new inputs, avoiding overfitting a resource to perform well just on specific test cases. When possible, input data should be made up of experimentally validated data. Indeed, similar results can make nearly impossible to establish a reliable ranking among participating tools making necessary to consider other aspects e.g. technical performance when similar scientific results are obtained.

### 4. Continuous release of data sets and assessment metrics

The periodic release of standardized benchmark data sets is crucial to allow independent training and testing. This is especially relevant for the refinement and development of new tools because it allows comparing these implementations with standardized data on already recognized tools. It is also relevant for developers because it allows to trace the evolution of a given resource — in terms of performance over time — for the same reference data and/or the performance of a given resource for an increasing number of reference data sets. Repositories such as Model Archive [Haas J. et al 2013], which aims to store protein models; UniProt [The UniProt Consortium 2017], which aims to provide a large-scale comprehensive and high-quality resource of protein sequences and functional annotations, and similar repositories facilitate data deposition and preservation. E-infrastructures for data handling like EUDAT [Lecarpentier D 2013] with specific services like B2Handle for providing persistent identifiers for data sets, B2Share and B2Drop for data storage, and B2Ffind for looking for data sets based on their metadata; or Zenodo can provide support and stable management for smaller projects. For continuous benchmarking efforts, regular standard test sets should be released and documented, for instance, in a short accompanying paper. Such continuous evolution fosters improvements in tools performance, and also aims at detecting and correcting software bugs as well as flaws in data sets used for benchmarking purposes. Moreover, substantial new features and functionalities could be discovered with the community support.

### 5. Regular Community Input

Regular meetings are envisioned with a large fraction of a given community present to discuss new developments and latest scoring methods, thereby driving the field to embrace new challenges and openly discuss best practice in assessing these aspects. For instance, in protein structure prediction, additional input of residue-residue contact predictions marked a game change in the field of *de novo* predictions. This success originated in community discussions at recurring meetings within CASP (CASP12, Dec 10-13 2016, Gaeta, Italy) and provides an excellent example of discussing new directions at community meetings. In addition to regular meetings, organizers should request feedback from the community after each event, for instance, via surveys and/or questionnaires to the participants. It is important to develop training and communication strategies to promote knowledge and experience exchanges within the community, and beyond. This is especially relevant for highly heterogeneous and geographically distributed communities. Importantly, results from a given benchmarking community might be of high relevance for other communities who are organized around downstream scientific aspects e.g. phylogenetic tree reconstruction methods highly depends on multiple sequence alignments quality; and/or focuses on the benchmarking of complex workflows scenarios rather than individual tools [Laurie S et al 2016].

## Specification and technical framework for online benchmarking

The constant growth of biological data represents unprecedented technical challenges as most software and data i) do not have, if any, structured descriptions; ii) lack useful metadata; iii) use non-standardized input and output formats; and iv) lack clear terms of use and/or licenses. Software source code, if available, is often obsolete, and makes it difficult to reproduce benchmark tests. Thus, it is necessary to develop a new technical framework to address, at least partially, these challenges. Such framework should implement approaches for managing, analyzing, and preserving both benchmarking data and metadata using standard formats for future reuse. The framework should, therefore, guarantee that data is **F**indable, **A**ccessible, **I**nteroperable, and **R**eusable according to the FAIR data principles [Wilkinson et al. 2016].

*OpenEBench* (Open ELIXIR Benchmarking and Technical Monitoring platform) is an attempt to address these challenges in a sustainable way under the ELIXIR-EXCELERATE umbrella. OpenEBench is an open infrastructure which focuses on supporting continuous automated community-driven benchmark efforts. OpenEBench aims at implementing different standards and best practice to facilitate the interoperability with other ELIXIR resources, including the ELIXIR Core data resources. OpenEBench also captures technical information about bioinformatics software aiming to provide a comprehensive description of scientific and technical performance of tools to developers, communities, end-users, and funders. The integration of different benchmark initiatives is meant to provide long term solution to the storage, analysis and publication of large scale heterogeneous data benchmarking ranging all across the biological data analysis challenges such as data mining, genomics, precision medicine, evolutionary and functional analysis. Finally its implementation has to be as inclusive as possible so it allows the integration of complex workflow, standalone software and web services.

### OpenEBench as an open benchmarking infrastructure

Community-driven benchmarking efforts can be: i) fully automated — which is the case for continuous online benchmarking; ii) manual — dependent on human expert predictors; or iii) hybrid — thus, combining both approaches like in CAFASP. We will focus on the first one giving the need to deal with enormous amounts of data and highly diverse scientific scenarios. To conduct continuous automated community-driven benchmarking challenges, it is necessary to have a stable infrastructure that allows significant and reliable participation. This infrastructure must scale according to the number of participants while allowing centralized collaborative efforts to define reference data sets. It should also implement a system for storing and sharing benchmarking results as well as performing benchmarking experiments. An extension to this infrastructure would allow complex bioinformatics workflows to be evaluated along with precise estimates of individual components and their fine-grained parameter tuning. Such a framework will optimally integrate the decision-making capabilities of communities and involved groups.

In order to engage and keep the interaction among community members, it is important to have a website which provides both friendly and unified programmatic access across different resources at the benchmarking platform. This central access point will facilitate data exchange, and promote results dissemination. To that end, service-oriented architectures should be designed using well-established web standards, such as those developed by the World Wide Web Consortium (W3C), the International Organization for Standardization (ISO) and the Internet Engineering Task Force (IETF^®^). Web-services should be platform-independent so they can be deployed and invoked from any platform and/or architecture. The early development of APIs (Application Programming interfaces) for the creation and deployment of web services will provide a way to conduct continuous automated benchmarking tests online. However, although web services provide the underlying basis for designing web architectures, further discussions about data, e.g. wide adoption of JSON format files as preferred interchange data format, and interoperability standards, e.g. implementation of FAIR data principles [Wilkinson et al. 2016], are needed.

The platform should implement mechanisms for guaranteeing reproducibility of conducted benchmarking experiments as well as to ensure the persistence of the involved data sets, e.g. input, output, results, and metrics. Besides, in order to avoid long execution cycles, it would be useful to develop specific testing and deployment paradigms that facilitate tools benchmarking. For instance, to deploy and execute different benchmarking experiments within the same technical environment, it is recommended the use of software container technologies, such as Docker [Boettiger 2015]. Software containers facilitate reproducibility, easy deployment and flexible building of collections of tools and search engines dedicated to specific scientific domains. Moreover, a conjunction of software containers could be used for benchmarking scientific workflows as well as to measure different metrics belonging to a specific benchmark. Thus, software orchestrators are needed to control those workflows. Two technologies will be deployed into OpenEBench: the widespread workflow manager Galaxy which offers a web-based user interface [Afgan et al. 2016], and Nextflow, a recently released framework which allows exact reproducibility and ease configuration and management of diverse tasks [Di Tommaso et al. 2017].

A schematic representation of the OpenEBench infrastructure can be found in Figure 6. As a highly interoperable infrastructure, it will make extensive use of APIs to communicate the central data warehouse with external providers, i.e. communities and software registries. Moreover, scientific benchmarking and technical monitoring results are accessible through an unified web-site. All data is also available to other web services via dedicated APIs.

**Figure 6.**
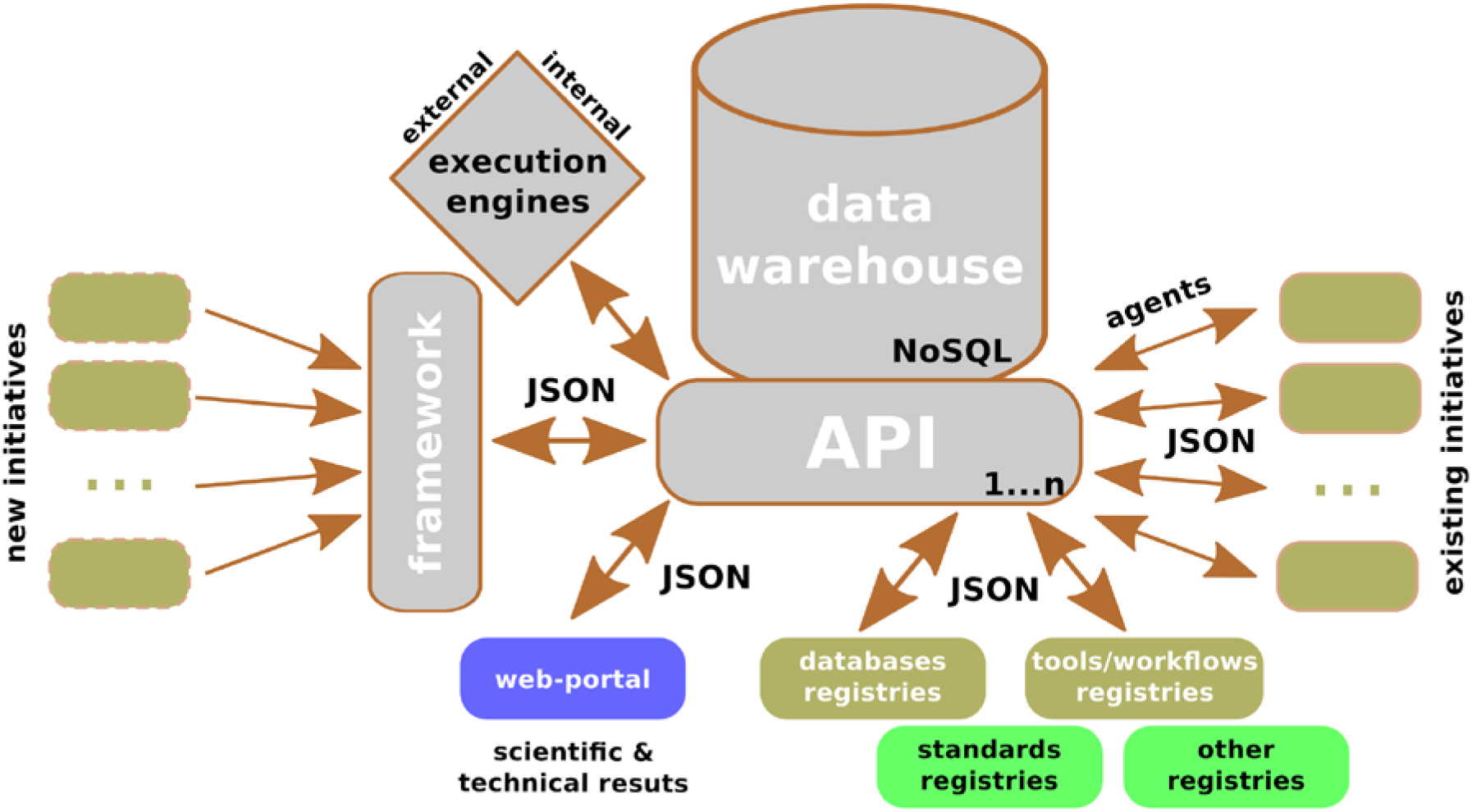
OpenEbench general overview representing input/output data sources (olive boxes), input data sources (lime boxes), output data resources (blue boxes), and technical components (gray boxes).

### Benchmark components and benchmarking process

In general, a benchmark can be defined by three main different components:

**Data resources**, including reference data sets for training and testing software resources and the ‘gold standard’ data set, if available. At the same time, the system should register participants contributions, preserving them in a centralized data warehouse available to the community (Figure 6]. Under some circumstances, the reference benchmark can be dynamic and updated through modalities agreed by the community (e.g. delayed structural publication for the CASP contest) so as to allow the evaluation of untrained tools.

**Guidelines and standards** for input and output formats allowing interoperability between computational resources. The benchmarking infrastructure will use standards for the design of the different services to be provided, as well as for the information transfer between them, both for data and metadata. Data standards would depend on the specific area under study, whilst metadata standards that define the semantics of the data are, generally, agnostics regarding the scientific domain e.g. ISO11179, which represents an example of an international standard for metadata-driven exchange of data in heterogeneous environments; ii) ISA-Tab, for complex metadata from ‘omics-based’ experiments [Sansone et al. 2012]; and iii) PDBx (PDB Exchange Dictionary), to define data content for deposition, annotation and archiving of PDB entries [Westbrook et al. 2005, Westbrook and Fitzgerald 2009].

**Scoring methodology and/or assessment tools** to compare the performance and rank the benchmark participants including scientific and technical metrics when possible.

Based on the aforementioned components, a standard continuous automated benchmarking service would consist of the following steps (Figure 7].

**Figure 7.**
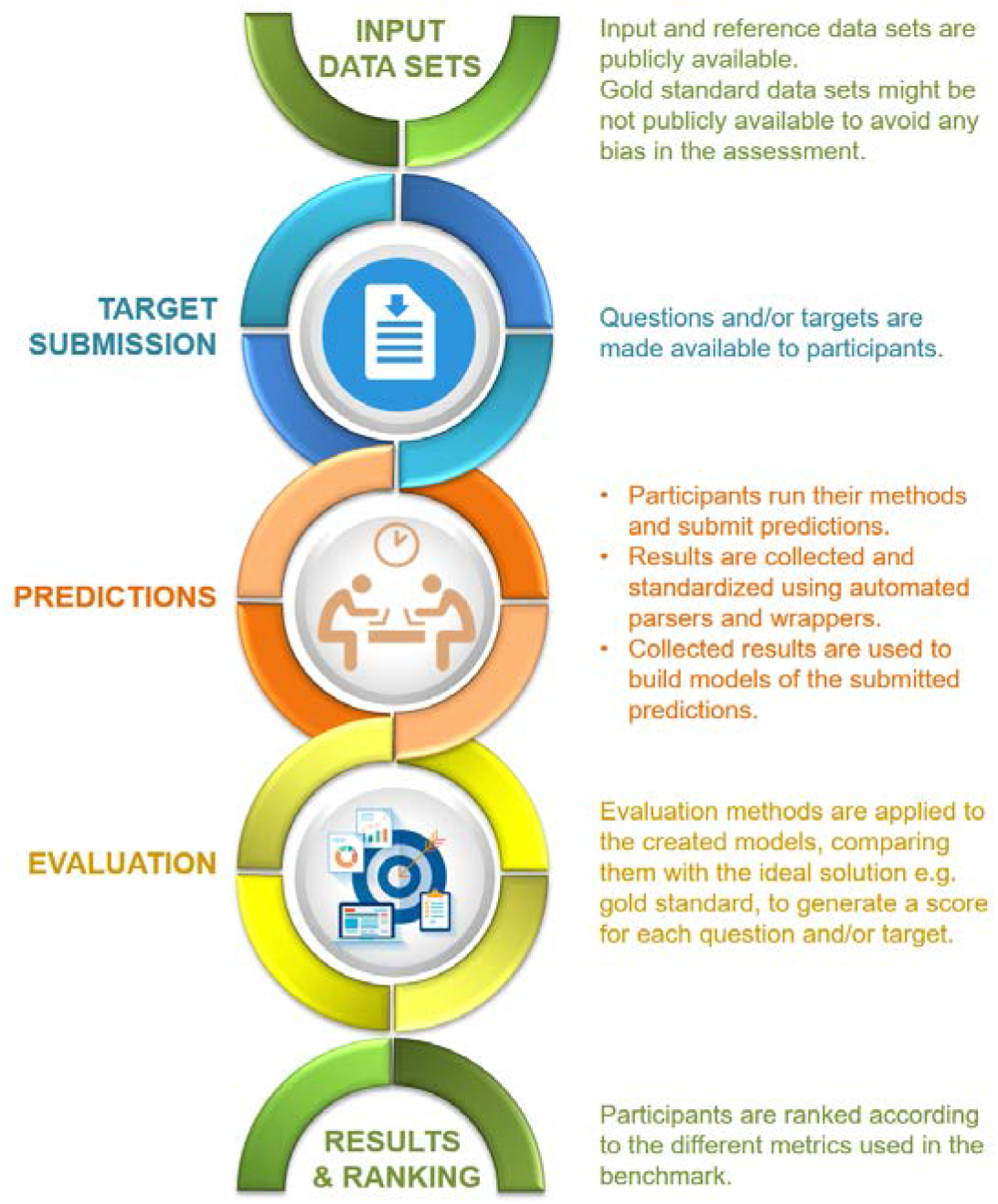
Stages of a benchmarking experiment within a scientific open challenge

### Interoperability and data formats

Interoperability of heterogeneous information systems is a key issue in bioinformatics [Thiam Yui et al. 2011, Sansone et al. 2012, Tenenbaum et al. 2014, Lopes and Oliveira 2015]. Within the context of collaborative challenges, interoperability is crucial to ensure that input and output data sets can be found, traced, analyzed, and shared inside and outside the community. The FAIR data principles [Wilkinson et al. 2016] can be accomplished by defining the proper data structures to model each piece of data using input and output standard formats. Ideally, each community should agree on data formats and common interfaces prior to the start of each benchmark initiative. Otherwise, data-related interoperability issues may arise when connecting different resources. One way to solve this situation is by using *ad-hoc*wrappers. In such scenario, it is important the role of communities in reaching consensus about how to interoperate in order to avoid decision-making based on factors that are not relevant to the research that is being conducted e.g. cost, availability, ease of use, etc. Lack of consensus may lead to numerous technical challenges for continuous automated benchmark efforts, for instance, how to integrate different data types e.g. functional, structural, sequences; and data formats; or how to facilitate rapid exchanges and access of heterogeneous data among disparate and distributed computational resources.

### Anonymizing data, secure access and participants visibility

A framework for hosting, processing, analyzing and sharing benchmark data requires a detailed bookkeeping system for tracking the server responses regarding i) who has accessed to what input data, and ii) who has submitted data to participate. Ideally, such framework also involves the use of an alerting system for identifying abnormal situations affecting specific participants and/or specific metrics in a given benchmark. Regarding data privacy and protection, when sensitive data is part of the challenge, either the submission of sensitive data must be anonymized, or both submission and reception should employ strong encryption algorithms, such as PGP (Pretty Good Privacy) [Zimmermann 1995] or AES (Advanced Encryption Standard) [Daemen and Rijmen 2001], and the appropriate data access policy should be enforced. Ideally, this decision should be agreed by the community members before starting the benchmark on how data should be processed. In addition, the benchmarking platform must allow users to participate at various visibility levels. The strictest would be “private”, where participants only can see their data in respect with other publicly available methods and no one else sees their data. The “community mode” allows all participants registered as members of the community to see each others data along with the public ones but data is not visible for users that do not participate in this community. Ideally, this data can be, at least partially, made publicly available once the benchmark campaign ends. The “submission mode” would allow participants to share their results with specific people e.g. reviewers when submitting a manuscript. An incentive to make data publicly available is to require participants to release at least one full submission in order to be part of any report and/or scientific publications derived from a given benchmark campaign. The default mode for any participating group must be “public” where results become publicly available once the infrastructure has automatically processed submitted data for a given benchmark edition. Publicly available data would be used to establish the reference results for any benchmark efforts and will be the only ones which are part of any report and/or scientific publication derived from community-driven benchmark efforts.

## Discussion and Conclusions

Current practices of self-evaluation of bioinformatics resources in terms of scientific and technical performance are usually limited to expert publications reviewing specific areas (not always exempt of biased) and periodic community challenges that evaluate tools on specific tasks, usually oriented to methods developers rather than general users. However, it is of high importance for developers and researchers being able to evaluate in the most continuous, open, automated and accessible way bioinformatics tools. Tools evaluation cannot be separated from the communities that develop the tools, since they are the ones with the expertise for identifying relevant scientific questions and technical aspects worth evaluating. Communities also drive the selection of appropriate strategies to measure answers to those questions as well as to identify the most suitable data sets to be used in the search of those answers. These efforts are of great value not only for tools developers to identify areas of improvement but importantly also for end-users that have to make informed choices of the best fitting resource for their scientific needs. It is also possible that funders will find this information useful for their evaluation of resources when deciding how to grant their resources.

The constant growth of available data and the vast amount of available bioinformatics software only reinforces the importance of such benchmarking efforts. In this scenario, large-scale efforts for developing, maintaining and extending centralized infrastructures which support those community efforts are essential. Leading initiatives like ELIXIR, EGI, EUDAT and NIH BD2k embrace those required platforms in order to secure standardization, guarantee interoperability, preserve reference data sets and strive to minimize the impact of budget and manpower constraints.

The quality and performance of existing benchmarking systems in bioinformatics have gradually improved over the years mainly due to the emergence of new scientific challenges and competitions after the success of initiatives like CASP [Moult et al. 1995] and BioCreAtIvE [Hirschman et al. 2005] as well as the lessons learnt from EVA [Rost and Eyrich 2001, Koh et al. 2003], LiveBench [Bujnicki et al. 2001, Rychlewski and Fischer 2005], and CAFA-SP [Fischer et al. 1999]. All together have fostered the creation of novel communities and initiatives, and their future potentials appear to be numerous and far-reaching [Costello and Stolovitzky 2013, Budge et al. 2015, Saez-Rodriguez et al. 2016].

As we pointed out before, essential in this endeavor is to determine what types of data should be collected for conducting assessments, what are the metrics to be measured, and which parameters would be used in a way that resources could be built around them. Making all these data and metadata interoperable is crucial to the development of bioinformatics tools with less bias and improved accuracy. Proper management of the metadata associated to any assessment i.e. evaluation environment, tools evaluated, parameters used; is important since it keep exact track on how tools have been evaluated. Under different conditions, it is possible that tools produce different results leading to different rankings. It is the task of the communities, assisted by systems like OpenEBench, to evaluate a relevant number of conditions opening up the possibility of evaluating further scenarios over time.

We strongly believe that most of bioinformatics disciplines would benefit from embracing community - driven benchmark efforts supported by an online centralized platform in parallel with the “classic” open challenges which are held face-to-face on scientific events such as competitions, hackathons and jamborees. In this sense, *OpenEBench* has been conceived to give support to that endeavor, providing means to collect, integrate and share data from different benchmarking efforts and communities. As our intention is to provide a dynamic resource, continuously available and updated, it will also provide the mechanisms to refine and expand certain data sets and metrics as new data become available in the future. With this in mind, we foresee *OpenEBench* functional assessment becoming a standard practice in the development and use of bioinformatics resources.

## Appendixes - tables

Table 1 - Summary of community-driven challenges

